# Oncolytic Newcastle disease virus promotes tumor cell apoptosis through SUMO-1-dependent PRKN-mediated degradation of c-Myc

**DOI:** 10.64898/2026.10.25.761721

**Authors:** Changrun Zhao, Ning Tang, Liya Shen, Jing Wang, Sonshu Meng, Jianghong Yu, Yiqian Zhu, Zunping Ke, Yang Qu, Lei Tan, Cuiping Song, Xusheng Qiu, Ying Liao, Yingjie Sun, Chan Ding

## Abstract

The oncogenic transcription factor c-Myc is aberrantly upregulated in many cancers and plays a central role in tumor cell proliferation and survival. However, direct pharmacological targeting of c-Myc remains challenging because of its intrinsically disordered structure. Newcastle disease virus (NDV) is an oncolytic virus with selective antitumor activity, but the molecular mechanisms by which NDV disrupts oncogenic signaling remain incompletely understood. Here, we show that NDV induces apoptosis in multiple tumor cell lines and suppresses tumor growth in an A549 xenograft model. NDV infection markedly reduced c-Myc protein levels in tumor cells in a time- and dose-dependent manner, and c-Myc knockdown further enhanced NDV-induced intrinsic apoptosis. Mechanistically, NDV promoted ubiquitin-proteasome-dependent degradation of c-Myc through ubiquitination at Lys142. Screening of candidate E3 ubiquitin ligases identified PRKN as a key mediator of NDV-induced c-Myc degradation. NDV infection induced SUMO-1-dependent nuclear translocation of PRKN, allowing PRKN to interact with nuclear c-Myc and promote its degradation. Disruption of this pathway by SUMO-1 or PRKN knockdown impaired PRKN nuclear accumulation, restored c-Myc stability, and attenuated NDV-induced apoptosis. Functionally, expression of the degradation-resistant c-Myc K142R mutant or restoration of c-Myc in c-Myc-knockdown cells protected tumor cells from NDV-induced apoptosis. Together, these findings identify a SUMO-1-PRKN-c-Myc signaling axis that mediates NDV-induced tumor cell apoptosis and reveal c-Myc destabilization as an important mechanism underlying the oncolytic activity of NDV.

**SIGNIFICANCE:** c-Myc is one of the most important oncogenic drivers in human cancer, but it remains difficult to target directly with conventional small-molecule inhibitors. This study identifies a previously unrecognized mechanism by which oncolytic Newcastle disease virus promotes c-Myc degradation. NDV induces SUMO-1-dependent nuclear translocation of the E3 ubiquitin ligase PRKN, which mediates c-Myc ubiquitination at Lys142 and promotes its proteasomal degradation. Loss of c-Myc activates intrinsic apoptotic signaling and enhances tumor cell apoptosis. These findings reveal a new NDV-regulated SUMO-1-PRKN-c-Myc axis and provide mechanistic insight into how oncolytic viruses can eliminate tumor cells by destabilizing key oncogenic proteins.

## INTRODUCTION

Cancer therapy remains limited by metastasis, drug resistance, treatment-related toxicity and tumor heterogeneity, which collectively reduce therapeutic efficacy and underscore the need for more effective and selective treatment strategies (1, 2). Oncolytic viruses (OVs) represent a promising therapeutic strategy because they preferentially infect and kill tumor cells while largely sparing normal tissues (3). Their antitumor effects are mediated not only by direct oncolysis but also by the induction of inflammatory and antitumor immune responses, thereby helping to overcome tumor-induced immunosuppression and immune evasion (3). Newcastle disease virus (NDV), a naturally occurring OV, has shown potent antitumor activity with low toxicity and has been reported to induce cell death in multiple tumor types (4–8). However, the molecular mechanisms underlying NDV-mediated tumor-selective killing remain incompletely defined, particularly regarding whether NDV acts by disrupting critical oncogenic pathways required for tumor cell survival.

c-Myc is a pleiotropic transcription factor that plays a central role in the regulation of cell growth, proliferation, metabolism and chromatin remodeling (9). It is also one of the four Yamanaka factors (OCT4, SOX2, KLF4 and c-MYC) required for reprogramming somatic cells into induced pluripotent stem cells (iPSCs) (10). In cancer biology, however, c-Myc is of particular interest because of its broad oncogenic functions (11–16). Aberrant activation or overexpression of c-Myc is frequently observed in human malignancies and is closely associated with tumor initiation, progression and poor prognosis(17, 18). In addition to driving malignant proliferation, c-Myc also contributes to immune evasion and metabolic reprogramming. For example, in non-small cell lung cancer, monocarboxylate transporter 1 (MCT1)-dependent acetate uptake promotes acetylation of c-Myc at Lys148, leading to its stabilization by ubiquitin-specific peptidase (USP) 10-mediated deubiquitination and resulting in enhanced programmed death-ligand 1 (PD-L1) expression, immune evasion and accelerated tumor cell proliferation (19). Because of its pivotal role in cancer, c-Myc has long been regarded as an attractive therapeutic target. However, direct targeting of c-Myc remains extremely challenging because of its intrinsically disordered and largely undruggable protein structure (9, 20). Therefore, alternative approaches that reduce c-Myc abundance or destabilize the c-Myc protein may provide new therapeutic opportunities for c-Myc-driven cancers.

Increasing evidence indicates that the abundance and transcriptional activity of c-Myc are tightly controlled by post-translational modifications, particularly phosphorylation, acetylation and ubiquitination (21). Among these, ubiquitination-dependent proteasomal degradation is a major mechanism governing c-Myc protein turnover. c-Myc is a short-lived protein, and its stability can be dynamically regulated by the ubiquitin-proteasome system through K48-linked polyubiquitination, which generally targets proteins for degradation (21, 22). Multiple E3 ubiquitin ligases and deubiquitinases have been reported to regulate c-Myc stability, thereby influencing tumor cell survival, proliferation and therapeutic response. For example, F-box and WD repeat domain-containing 7 (FBW7) is a well-characterized E3 ligase that promotes c-Myc ubiquitination and degradation in a phosphorylation-dependent manner (23, 24), whereas USP28 and USP36 antagonize c-Myc degradation and enhance its stability (25, 26). In addition, dysregulation of the ubiquitin-proteasome system is increasingly recognized as an important mechanism contributing to persistent c-Myc activation in cancer (21). These findings suggest that manipulating the post-translational stability of c-Myc may represent a feasible strategy to overcome the limitations of direct c-Myc inhibition.

Because viruses frequently reprogram host protein homeostasis and ubiquitin-mediated signaling to regulate intracellular pathways and cell fate (27, 28), it is plausible that NDV may exert part of its antitumor activity by modulating the stability of oncogenic proteins such as c-Myc. NDV has also shown considerable promise as both an oncolytic agent and a genetic engineering vector, and recombinant NDV strains have demonstrated enhanced antitumor efficacy in preclinical and clinical studies (29, 30). Nevertheless, whether NDV suppresses tumor growth by inducing ubiquitin-dependent degradation of c-Myc remains unknown. In this study, we demonstrate that NDV induces apoptosis in multiple tumor cell types and suppresses tumor growth by promoting ubiquitin-proteasome-dependent degradation of c-Myc. Mechanistically, NDV triggers polyubiquitination of c-Myc at Lys142, leading to its destabilization and degradation. We further show that NDV promotes SUMO1-dependent nuclear translocation of the E3 ubiquitin ligase PRKN, which mediates c-Myc ubiquitination and is required for NDV-induced tumor cell apoptosis. These findings uncover a previously unrecognized NDV-SUMO1-PRKN-c-Myc signaling axis and provide new mechanistic insight into the oncolytic action of NDV.

## RESULTS

### NDV induces tumor cell apoptosis in vitro and in vivo

NDV has been reported to induce apoptosis in a variety of cancer cells (4, 6, 8). To further assess the pro-apoptotic effect of NDV, we used the NDV Herts/33 strain (hereafter referred to as NDV) to examine its activity in lung adenocarcinoma cells (A549), cervical cancer cells (HeLa), pancreatic cancer cells (PANC-1), colon cancer cells (HT-29) and breast cancer cells (MCF-7). Annexin V/PI staining showed that NDV infection induced apoptosis to different extents in different cell lines (Fig. 1A-E). Consistently, immunoblot analysis showed cleavage of caspase-3 and PARP, two major biochemical markers of apoptosis, in lung, cervical, breast, pancreatic and colon cancer cells at 24 h post-infection (hpi) (Fig. 1F-J), although the extent of cleavage varied among cell lines.

**Figure 1.**
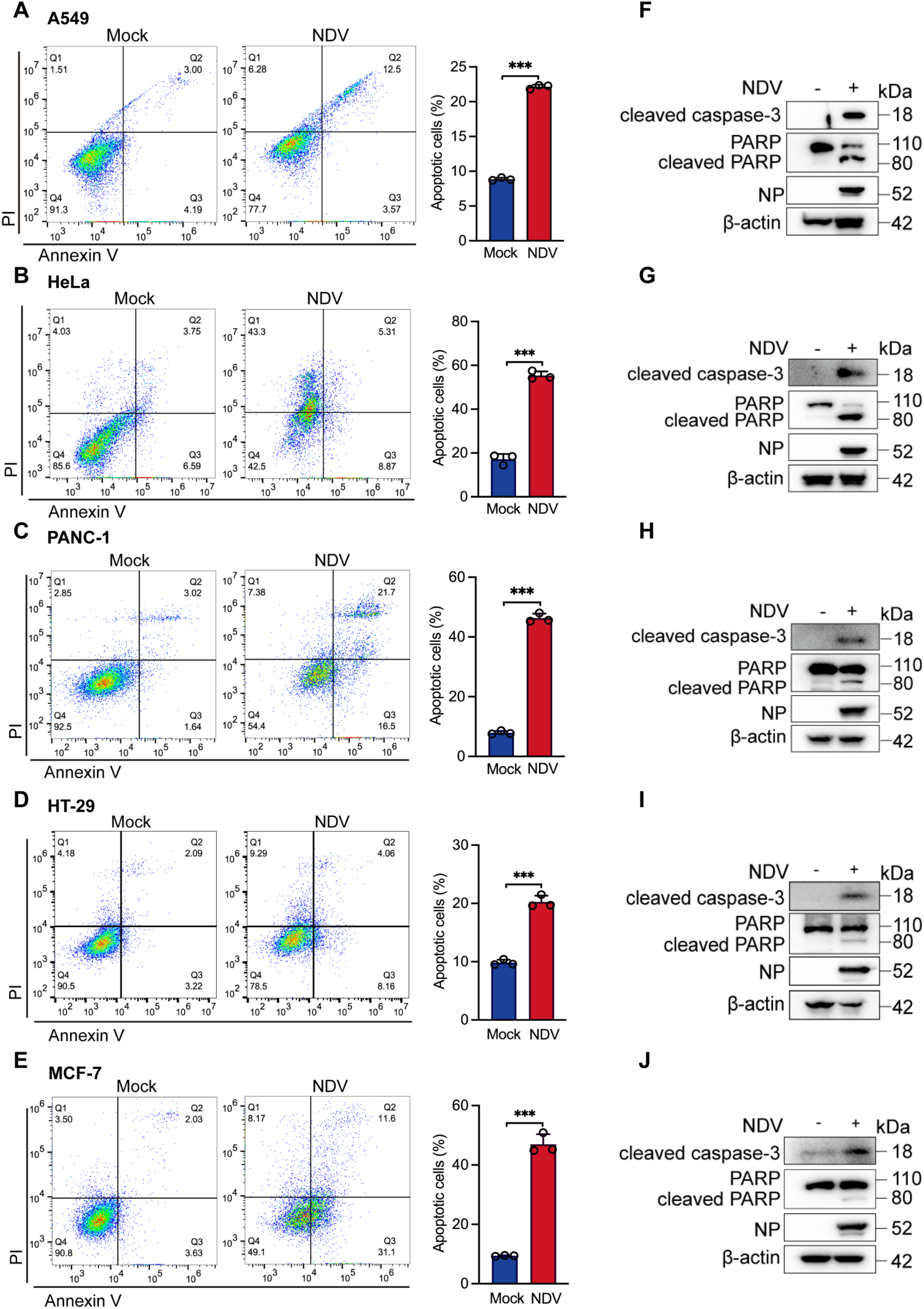
NDV induces apoptosis in multiple tumor cell lines. (A-E) A549 non-small cell lung cancer cells (A), HeLa cervical cancer cells (B), PANC-1 pancreatic cancer cells (C), HT-29 colon cancer cells (D), and MCF-7 breast cancer cells (E) were mock-treated or infected with NDV at an MOI of 1. Apoptosis was analyzed by flow cytometry at 24 hpi, and the percentages of apoptotic cells are shown. (F-J) Western blot analysis of apoptosis-related proteins in A549 (F), HeLa (G), PANC-1 (H), HT-29 (I), and MCF-7 (J) cells following NDV infection. Cleaved caspase-3, PARP, and cleaved PARP were detected, with β-actin used as the loading control and NDV NP used as a marker of infection. Data are presented as means from three independent experiments. ***P < 0.001.

We next evaluated the antitumor activity of NDV in vivo using an A549 xenograft model (Fig. 2A). NDV treatment had no significant effect on body weight (Fig. 2B), but markedly suppressed tumor growth (Fig. 2C and D). Increased apoptosis in NDV-treated tumors was confirmed by TUNEL staining and elevated cleaved caspase-3 expression (Fig. 2E), and further supported by increased cleaved caspase-9 and cleaved caspase-3 levels by western blotting (Fig. 2F). Together, these results indicate that NDV activates apoptotic signaling and induces tumor cell apoptosis both *in vitro* and *in vivo*.

**Figure 2.**
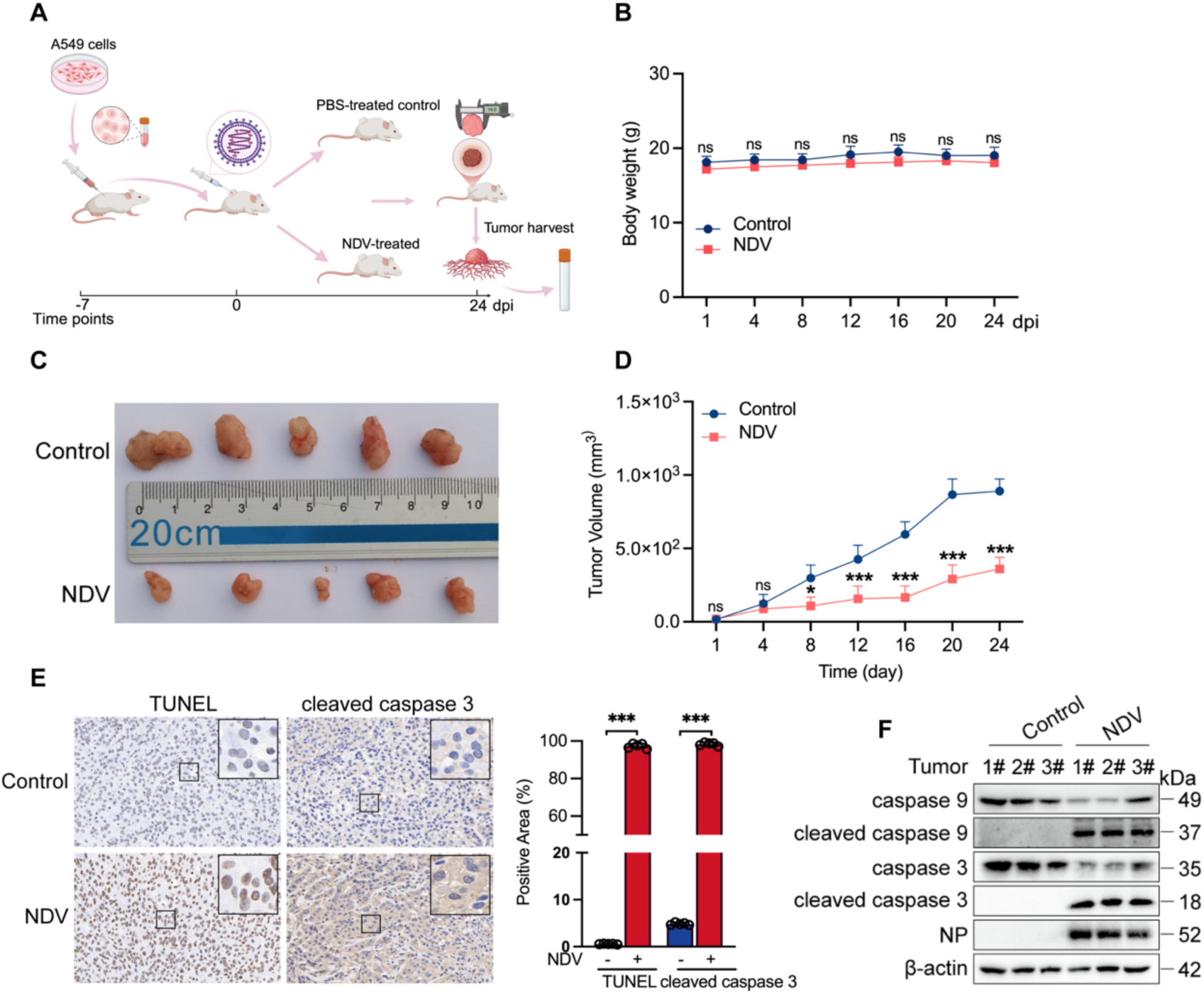
NDV suppresses tumor growth in vivo by inducing apoptosis. (A) Schematic illustration of the A549 xenograft model and NDV treatment schedule. A549 cells were subcutaneously injected into mice to establish tumor-bearing models. Seven days after subcutaneous injection, tumor-bearing mice were randomized and treated intravenously with NDV, 10^8.5^ EID_50_/100 μL, or PBS twice weekly for 24 days. (B) Body weight curves of A549 tumor-bearing mice during NDV treatment, n = 5. (C and D) Representative tumor images and tumor growth curves showing the inhibitory effect of NDV on A549 xenograft tumor growth, n = 5. (E) TUNEL and cleaved caspase-3 staining of tumor sections, with quantification of positive staining areas from five randomly selected fields using ImageJ. (F) Western blot analysis of caspase-9, cleaved caspase-9, caspase-3, and cleaved caspase-3 in tumor tissues. β-actin was used as the loading control, and NDV nucleoprotein (NP) was used as a marker of viral infection. Data are presented as means from three independent experiments. *P < 0.05, ***P < 0.001; ns, not significant.

### NDV induces the degradation of c-Myc in tumor cells

c-MYC, SOX2, KLF4 and OCT4 are four core transcription factors involved in somatic cell reprogramming and are also closely associated with tumor development. Analysis of The Cancer Genome Atlas (TCGA) and Human Protein Atla (HPA) datasets showed that c-MYC, SOX2, KLF4 and OCT4 are broadly expressed in human malignancies, with c-MYC displaying particularly prominent upregulation (Fig. 3A-D). We therefore examined whether NDV affects the expression of these reprogramming-associated transcription factors. In A549 cells, NDV infection markedly reduced endogenous c-Myc and KLF4 protein levels at 18 and 24 hpi, whereas SOX2 and OCT4 were increased (Fig. 3E). Consistent changes were also detected following infection with the mesogenic strain Mukteswar and the lentogenic strain La Sota (Fig. 3F and G), indicating that the effect observed with the velogenic Herts/33 strain is shared by NDV strains of different pathotypes. However, UV-inactivated NDV failed to induce these changes, suggesting that active viral infection is required (Fig. 3H). We next examined whether these alterations also occur *in vivo*. Similar changes in SOX2, KLF4 and OCT4 expression were observed in NDV-treated xenograft tumors (Fig. 3I), consistent with the *in vitro* results (Fig. 3E-G). Importantly, c-Myc expression was markedly reduced in NDV-treated tumors, as confirmed by immunohistochemistry (Fig. 3J). Together, these findings suggest that NDV broadly modulates reprogramming-associated transcription factors both *in vitro* and *in vivo*. Among them, the marked downregulation of c-Myc prompted us to further investigate its potential role in NDV-mediated apoptosis and tumor suppression.

**Figure 3.**
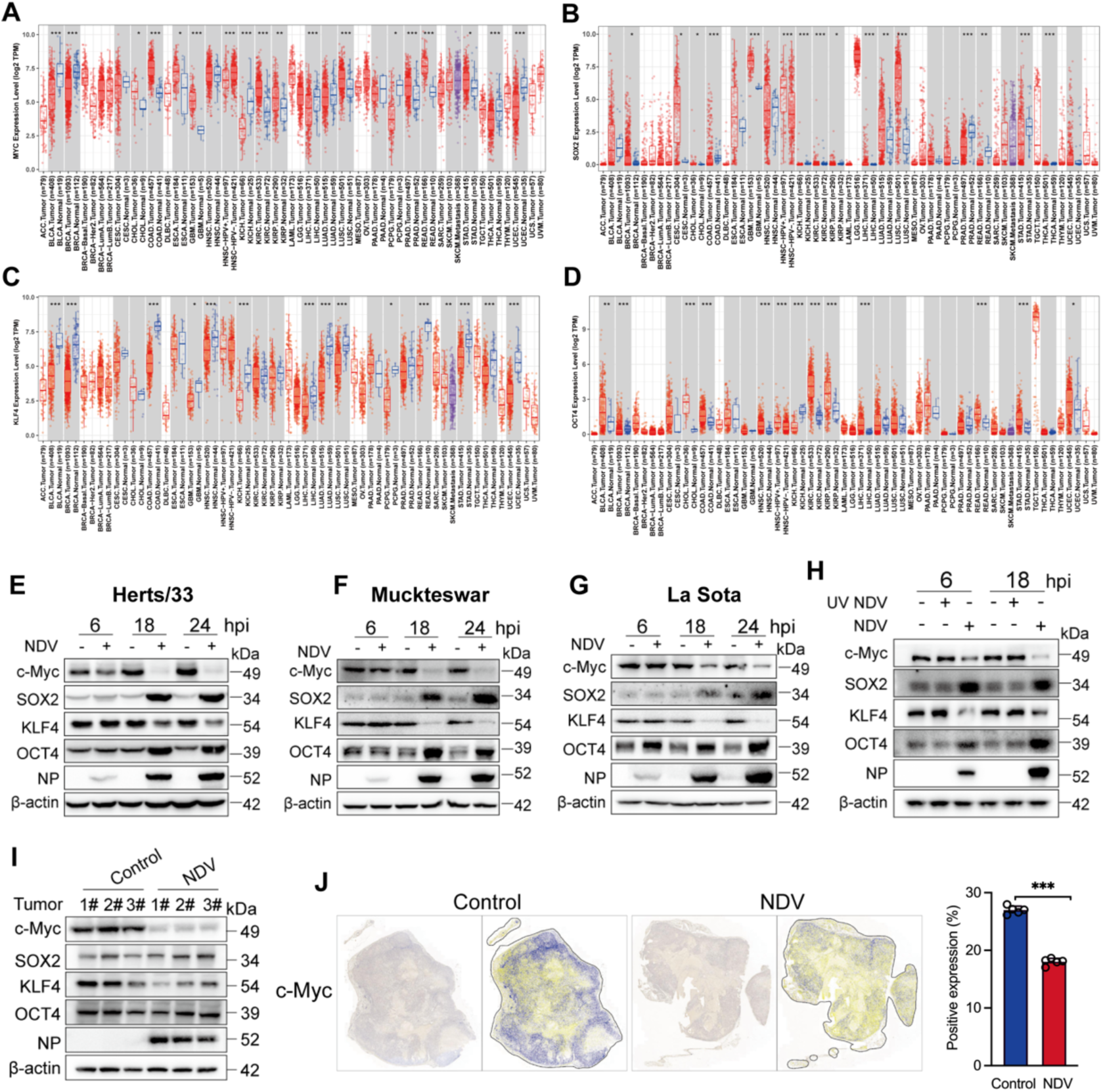
NDV downregulates c-Myc in tumor cells and xenograft tumors. (A-D) Expression profiles of c-Myc (A), SOX2(B), KLF4(C), and OCT4(D) in normal and tumor tissues were analyzed using The Cancer Genome Atlas (TCGA) and Human Protein Atlas (HPA) databases. (E-G) Western blot analysis of c-Myc, SOX2, KLF4, and OCT4 protein levels in A549 cells infected with different NDV strains, including Herts/33 (E), Mukteswar (F), and La Sota (G), at an MOI of 1 for 6, 18, and 24 h. (H) Western blot analysis of c-Myc, SOX2, KLF4, and OCT4 protein levels in A549 cells infected with live NDV or UV-inactivated NDV at an MOI of 1 for 6 and 18 h. (I) Western blot analysis of caspase-9, cleaved caspase-9, caspase-3, and cleaved caspase-3 in A549 xenograft tumor tissues. β-actin was used as the loading control, and NDV nucleoprotein (NP) was used as a marker of viral infection. (J) Immunohistochemical staining of c-Myc in A549 xenograft tumor tissues, with quantification of positive staining areas using ImageJ. Data are presented as means from three independent experiments. ***P < 0.001.

### NDV-induced c-Myc downregulation promotes intrinsic apoptosis in tumor cells

c-Myc is highly expressed in multiple tumor types and promotes tumor cell proliferation (17, 18). Given the central oncogenic role of c-Myc, we next focused on this factor. NDV induced c-Myc degradation in multiple tumor cell lines, including HeLa, MCF-7, PANC-1 and HT29 (Fig. 4A-D). This effect was also dose-dependent across all three NDV strains tested (Fig. 4E-G). Consistently, immunofluorescence analysis showed that c-Myc was predominantly localized in the nucleus of A549 cells, but its signal was markedly reduced at 18 and 24 hpi following NDV infection (Fig. 4H). These results indicate that NDV downregulates c-Myc protein levels in tumor cells in a time- and dose-dependent manner.

**Figure 4.**
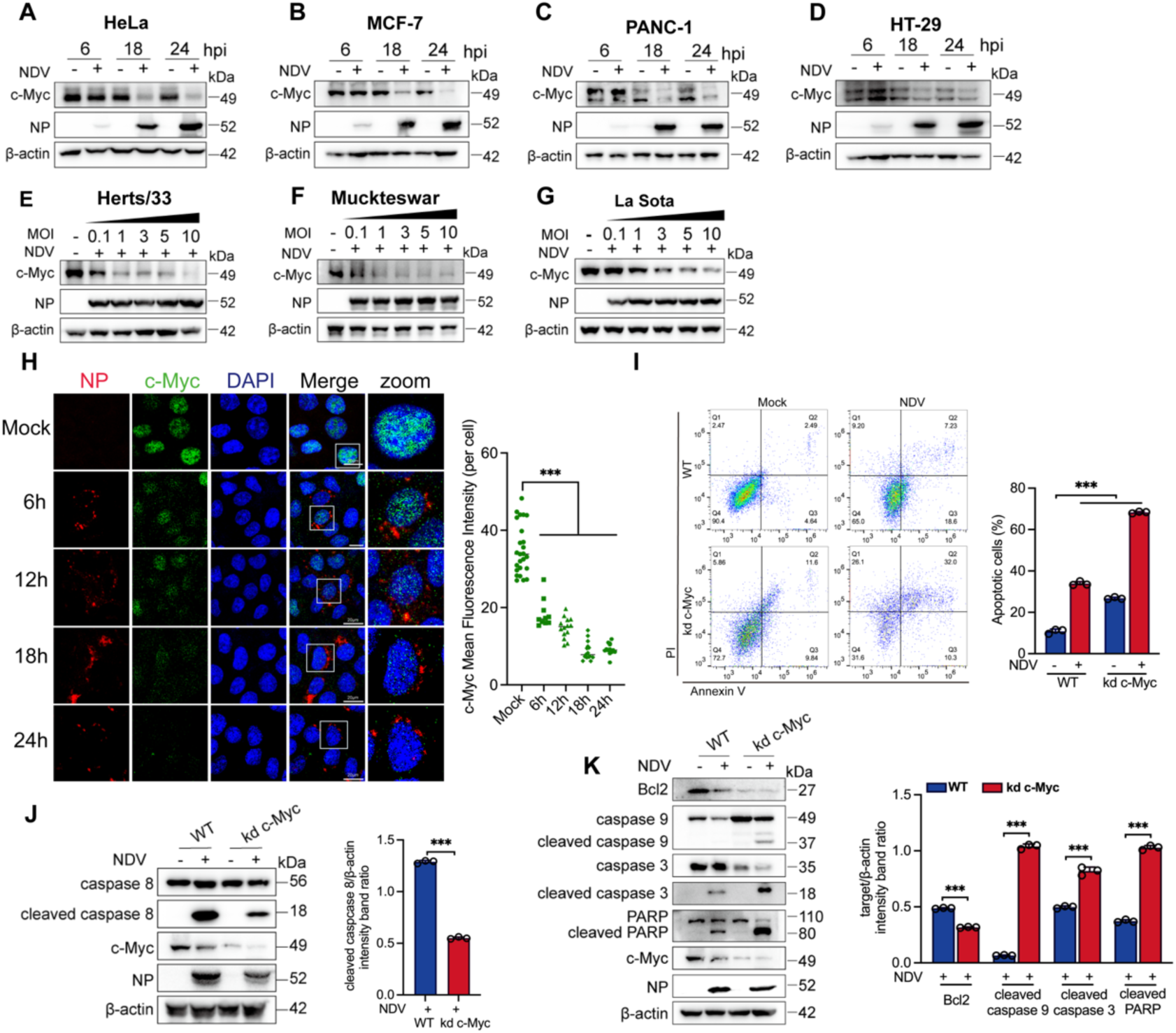
NDV-induced c-Myc downregulation promotes tumor cell apoptosis. (A-D) Western blot analysis of c-Myc protein levels in HeLa (A), MCF-7 (B), PANC-1 (C), and HT-29 (D) cells infected with NDV at an MOI of 1 for 6, 18, and 24 h. (E-G) Western blot analysis of c-Myc protein levels in A549 cells infected with different doses of NDV Herts/33 (E), Mukteswar (F), or La Sota (G) strains at MOIs of 0.1, 1, 3, 5, and 10 for 18 h. (H) Immunofluorescence analysis of c-Myc and NDV NP in A549 cells mock-treated or infected with NDV at an MOI of 1 for 6, 12, 18, and 24 h. Mean fluorescence intensity of c-Myc per cell was quantified using ImageJ. (I) Flow cytometric analysis of apoptosis in wild-type and c-Myc-knockdown A549 cells after mock treatment or NDV infection at an MOI of 1 for 24 h, and the percentages of apoptotic cells are shown. (J) Western blot analysis of caspase-8, cleaved caspase-8, and c-Myc in wild-type and c-Myc-knockdown A549 cells after mock treatment or NDV infection. β-actin was used as the loading control, and NDV NP was used as a marker of viral infection. Densitometric quantification of cleaved caspase-8 normalized to β-actin is shown. (K) Western blot analysis of Bcl-2, caspase-9, cleaved caspase-9, caspase-3, cleaved caspase-3, PARP, cleaved PARP, and c-Myc in wild-type and c-Myc-knockdown A549 cells after mock treatment or NDV infection. β-actin was used as the loading control, and NDV NP was used as a marker of viral infection. Densitometric quantification of Bcl-2, cleaved caspase-9, cleaved caspase-3, and cleaved PARP normalized to β-actin is shown. Data are presented as means from three independent experiments. ***P < 0.001.

To examine whether c-Myc reduction contributes to NDV-induced oncolysis, we silenced c-Myc in tumor cells. c-Myc knockdown induced apoptosis and further enhanced NDV-induced apoptosis (Fig. 4I). Under these conditions, cleaved caspase-8 was reduced (Fig. 4J), whereas cleaved caspase-9 and cleaved caspase-3 were increased (Fig. 4K), indicating preferential activation of the intrinsic apoptotic pathway. Collectively, these findings suggest that NDV-mediated c-Myc downregulation contributes to tumor cell apoptosis, and that loss of c-Myc sensitizes tumor cells to NDV-induced intrinsic apoptosis.

### NDV induces proteasomal degradation of c-Myc through ubiquitination at K142

The ubiquitin-proteasome system and autophagy-lysosome pathway are the two major mechanisms of intracellular protein degradation. To identify the pathway responsible for NDV-induced c-Myc degradation, cells were treated with MG132 or chloroquine (CQ). MG132, but not CQ, restored c-Myc protein levels following NDV infection (Fig. 5A and B), indicating that NDV-induced c-Myc degradation is mainly mediated by the ubiquitin-proteasome pathway. Consistently, co-immunoprecipitation analysis showed that NDV infection markedly increased c-Myc ubiquitination (Fig. 5C). To map the region responsible for NDV-induced c-Myc degradation, we generated two c-Myc truncation mutants, 1-300 aa and 300-454 aa (Fig. 5D). NDV induced degradation of the 1-300 aa fragment, indicating that the critical region lies within the N-terminal portion of c-Myc (Fig. 5E). This region contains nine lysine residues, which were individually mutated to arginine. Among these mutants, only K142R markedly impaired NDV-induced c-Myc ubiquitination, whereas the other lysine mutations had little effect (Fig. 5F). These results indicate that NDV promotes proteasomal degradation of c-Myc through ubiquitination at K142, identifying K142 as a key residue controlling c-Myc stability.

**Figure 5.**
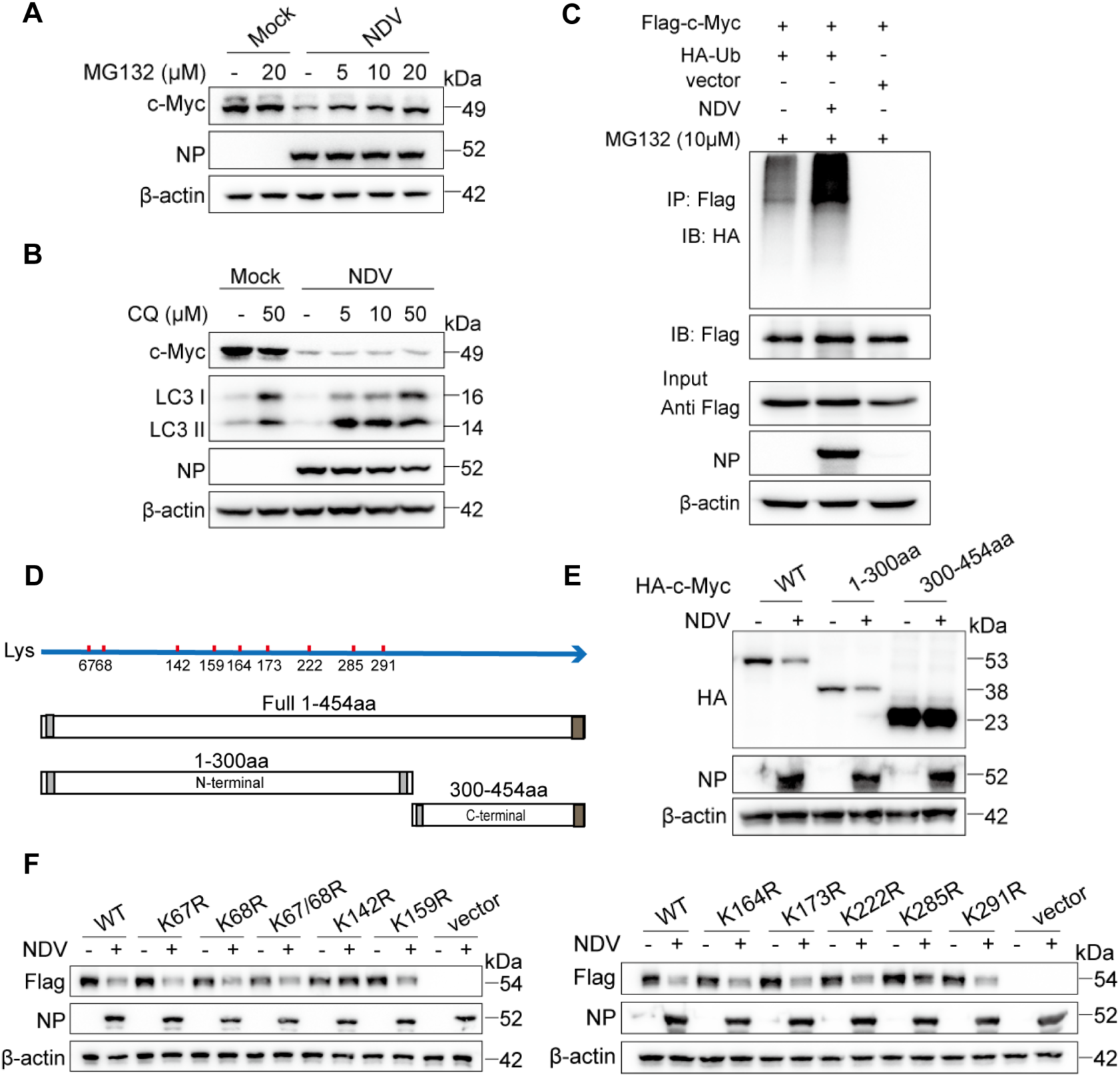
NDV promotes proteasomal degradation of c-Myc through ubiquitination at Lys142. (A and B) A549 cells were infected with NDV at an MOI of 1 for 18 h in the presence or absence of MG132 (A) or CQ (B). c-Myc protein levels were analyzed by western blot. (C) HEK-293T cells were co-transfected with Flag-c-Myc and HA-ubiquitin for 24 h, and then infected with NDV in the presence of MG132. At 18 hpi, Flag-c-Myc was immunoprecipitated with an anti-Flag antibody, and c-Myc ubiquitination was analyzed by western blot. (D) Schematic diagram of HA-tagged full-length and truncated c-Myc constructs, with potential ubiquitination sites indicated. (E) A549 cells were transfected with HA-tagged full-length c-Myc or truncated c-Myc constructs, including aa 1 to 300 and aa 300 to 454. At 24 h post transfection, cells were infected with NDV at an MOI of 1 and harvested at 18 hpi for western blot analysis. (F) A549 cells were transfected with Flag-tagged c-Myc lysine-to-arginine mutants, including K67R, K68R, K67/68R, K142R, K159R, K164R, K173R, K222R, K285R, and K291R. At 24 h post transfection, cells were infected with NDV at an MOI of 1 and harvested at 18 hpi. Flag-c-Myc protein levels were analyzed by western blot.

### NDV promotes c-Myc degradation through SUMO-1-dependent nuclear translocation of PRKN

After identifying Lys142 as a critical site for c-Myc ubiquitination and degradation, we next sought to identify the host E3 ligase responsible for this process. Based on previous reports and prediction using UbiBrowser v3.0, seven candidate E3 ligases, including TRPC4AP, STUB1, FBXW7, RNF187, PRKN, FBXW11, and SKP2, were selected for further analysis. The roles of these candidate E3 ligases in NDV-induced c-Myc degradation were assessed using specific siRNAs. PRKN knockdown markedly attenuated NDV-induced c-Myc degradation (Fig. 6A), with knockdown efficiency confirmed by RT-qPCR (Fig. 6B). Co-immunoprecipitation analysis further showed an interaction between PRKN and c-Myc, supporting the involvement of PRKN in c-Myc degradation during NDV infection (Fig. 6C).

**Figure 6.**
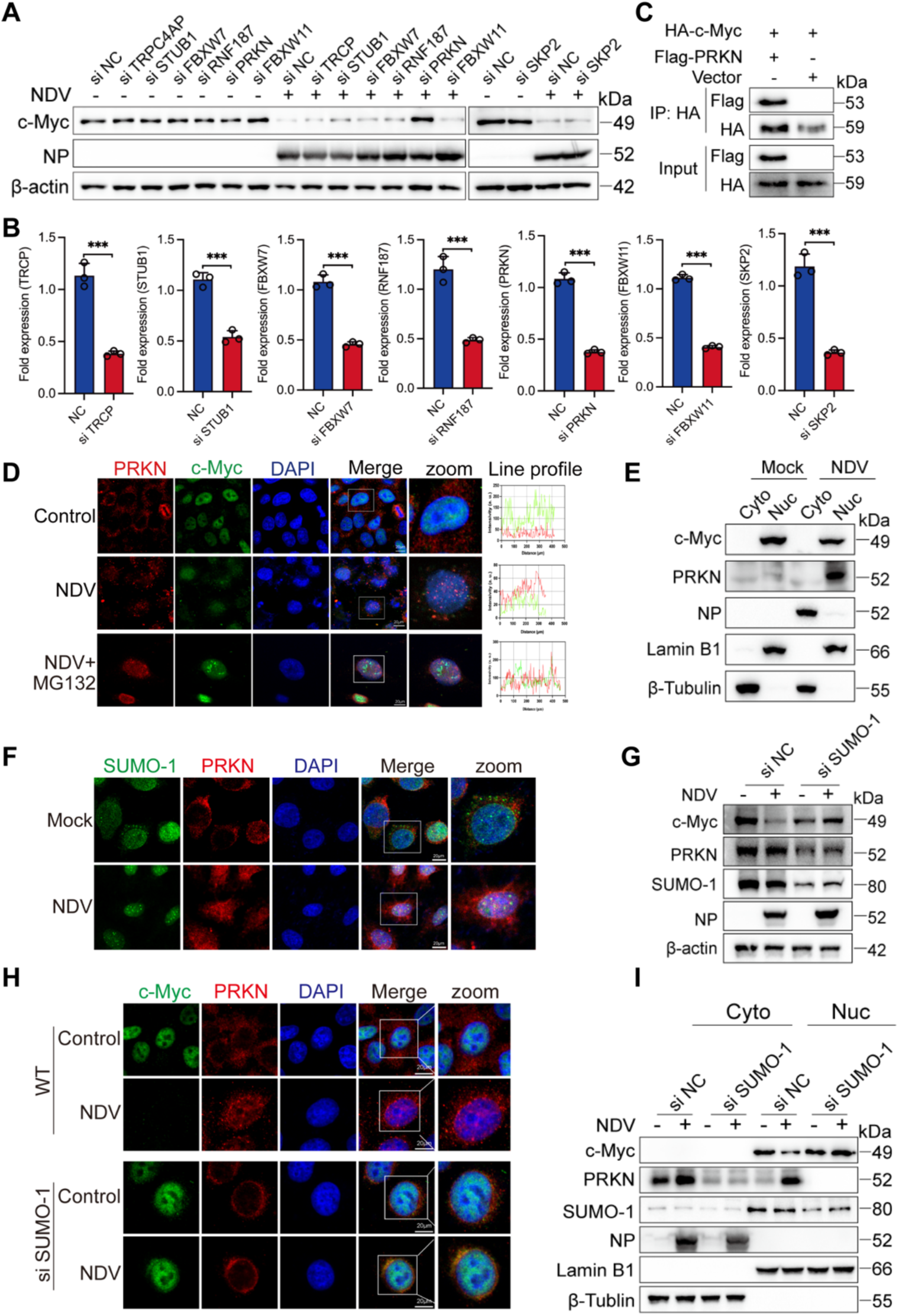
NDV induces SUMO-1-dependent nuclear translocation of PRKN to promote c-Myc degradation. (A and B) A549 cells were transfected with scrambled control siRNA or siRNAs targeting TRPC4AP, STUB1, FBXW7, RNF187, PRKN, FBXW11, or SKP2 for 24 h, followed by NDV infection at an MOI of 1 for 18 h. c-Myc protein levels were analyzed by western blot (A), and knockdown efficiency was confirmed by RT-qPCR (B). (C) HEK-293T cells were co-transfected with HA-c-Myc and Flag-PRKN for 24 h, followed by NDV infection at an MOI of 1 for 18 h. HA-c-Myc was immunoprecipitated with an anti-HA antibody, and the interaction between c-Myc and PRKN was analyzed by western blot. (D) A549 cells were mock-treated or infected with NDV at an MOI of 1 for 18 h in the presence or absence of MG132, followed by immunofluorescence analysis of PRKN and c-Myc. Line-scan intensity profiles of PRKN and c-Myc signals across the indicated regions of interest are shown. (E) A549 cells were mock-treated or infected with NDV at an MOI of 1 for 18 h, followed by subcellular fractionation. c-Myc and PRKN levels in the cytoplasmic and nuclear fractions were analyzed by western blot. (F) A549 cells were mock-treated or infected with NDV at an MOI of 1 for 18 h, followed by immunofluorescence analysis of SUMO-1 and PRKN. (G) A549 cells were transfected with scrambled control siRNA or SUMO-1-specific siRNA for 24 h, followed by NDV infection at an MOI of 1 for 18 h. Total c-Myc, PRKN, and SUMO-1 protein levels were analyzed by western blot. (H) A549 cells were transfected with scrambled control siRNA or SUMO-1-specific siRNA for 24 h, followed by NDV infection at an MOI of 1 for 18 h. c-Myc and PRKN localization was analyzed by immunofluorescence. (I) A549 cells were transfected with scrambled control siRNA or SUMO-1-specific siRNA for 24 h, followed by NDV infection at an MOI of 1 for 18 h and subcellular fractionation. c-Myc and PRKN levels in the cytoplasmic and nuclear fractions were analyzed by western blot. Data are presented as means from three independent experiments. ***P < 0.001.

Because c-Myc is mainly localized in the nucleus whereas PRKN is predominantly cytoplasmic under basal conditions, the subcellular localization of PRKN during NDV infection was examined. Immunofluorescence analysis showed that PRKN was mainly distributed in the cytoplasm of uninfected cells but accumulated as punctate signals in the nucleus after NDV infection. In the presence of MG132, nuclear PRKN partially colocalized with c-Myc, suggesting that PRKN may target c-Myc for proteasomal degradation in the nucleus (Fig. 6D). Consistently, subcellular fractionation confirmed that endogenous PRKN was markedly increased in the nuclear fraction following NDV infection (Fig. 6E). Previous studies have reported that SUMO-1 modification regulates the subcellular localization and autoubiquitination activity of PRKN (31). Immunofluorescence analysis showed that NDV infection induced nuclear accumulation of SUMO-1, accompanied by PRKN aggregation in the nucleus (Fig. 6F). SUMO-1 knockdown also blocked NDV-induced c-Myc degradation, supporting the requirement of SUMO-1 for PRKN-mediated c-Myc degradation (Fig. 6G). Mechanistically, SUMO-1 knockdown impaired NDV-induced nuclear accumulation of PRKN, as shown by immunofluorescence analysis (Fig. 6H). Consistently, subcellular fractionation confirmed reduced PRKN accumulation in the nuclear fraction after SUMO-1 knockdown (Fig. 6I). Collectively, these findings demonstrate that NDV promotes c-Myc degradation by inducing SUMO-1-dependent nuclear translocation of PRKN.

### c-Myc degradation is critical for NDV-induced tumor cell apoptosis

Having established that NDV promotes c-Myc ubiquitination and degradation at Lys142, the functional significance of this event in NDV-mediated antitumor activity was further examined. Cells stably expressing the degradation-resistant c-Myc K142R mutant showed reduced sensitivity to NDV-induced apoptosis (Fig. 7A). Consistently, c-Myc K142R expression increased Bcl-2 levels and decreased the activation of caspase-9 and caspase-3 after NDV infection (Fig. 7B), indicating that c-Myc degradation at Lys142 contributes to NDV-induced tumor cell apoptosis. Similarly, PRKN knockdown significantly attenuated NDV-induced apoptosis (Fig. 7C), accompanied by increased Bcl-2 expression and reduced levels of cleaved caspase-9 and cleaved caspase-3 (Fig. 7D). To further assess the role of c-Myc in NDV-mediated antitumor activity, wild-type c-Myc was reconstituted in c-Myc-knockdown cells, referred to as re-c-Myc cells. Restoration of c-Myc promoted tumor cell proliferation and reduced NDV-induced apoptosis (Fig. 7E). In line with this effect, Bcl-2 expression was increased, whereas cleaved caspase-9 and cleaved caspase-3 levels were decreased in re-c-Myc cells following NDV infection (Fig. 7F). Notably, reconstitution of the degradation-resistant c-Myc K142R mutant provided stronger protection against NDV-induced apoptosis than wild-type c-Myc (Fig. 7G), accompanied by increased Bcl-2 expression and decreased activation of caspase-9 and caspase-3 (Fig. 7H). Together, these findings demonstrate that NDV promotes tumor cell apoptosis by targeting c-Myc for PRKN-mediated degradation at Lys142.

**Figure 7.**
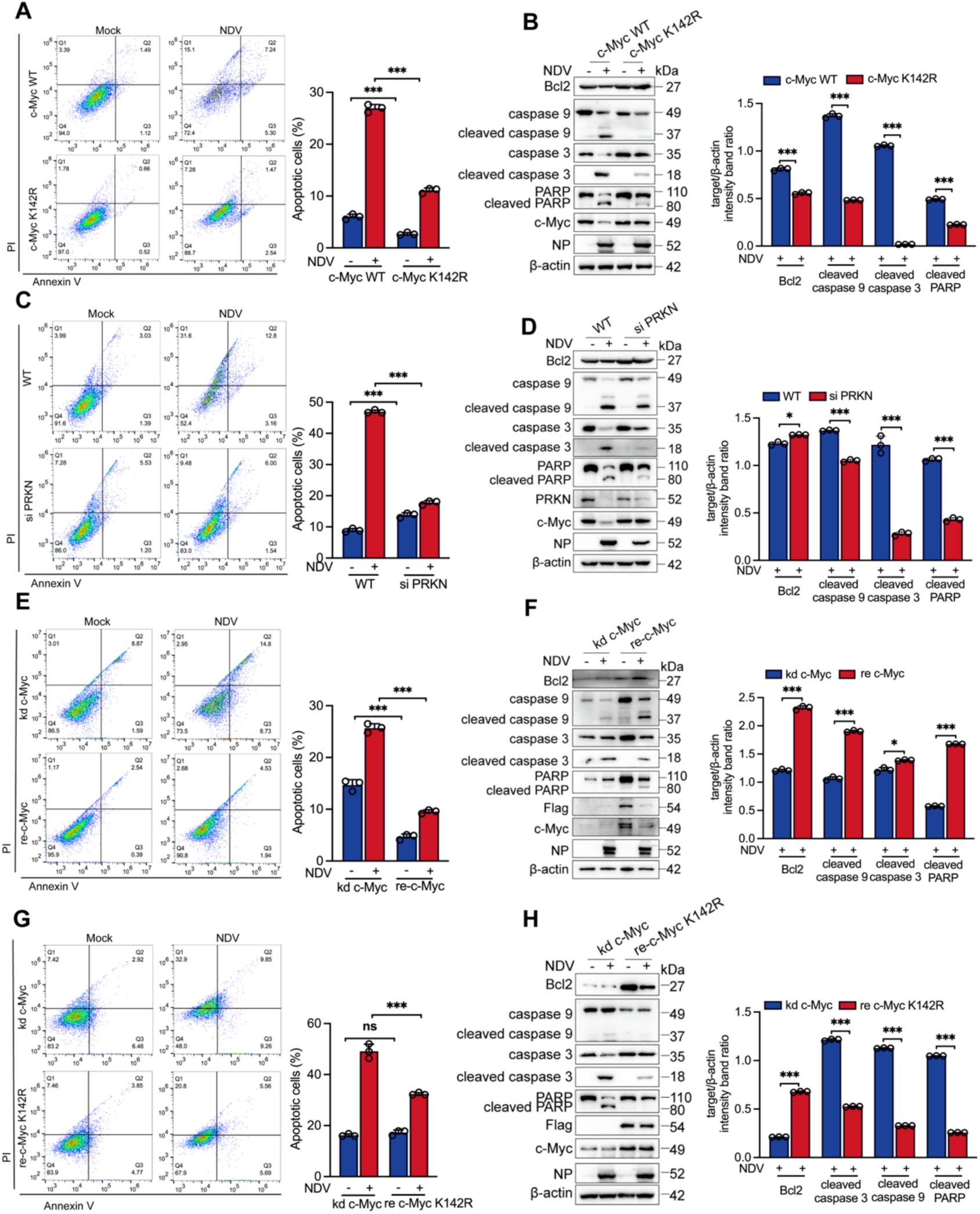
NDV promotes tumor cell apoptosis by targeting c-Myc. (A and B) A549 cells stably expressing wild-type c-Myc or the degradation-resistant c-Myc K142R mutant were mock-treated or infected with NDV at an MOI of 1 for 24 h. Apoptosis was analyzed by flow cytometry, and the percentages of apoptotic cells are shown (A). Apoptosis-related protein levels were analyzed by western blot, with densitometric quantification relative to β-actin shown (B). (C and D) A549 cells were transfected with scrambled control siRNA or PRKN-specific siRNA for 24 h, followed by NDV infection at an MOI of 1 for 24 h. Apoptosis was analyzed by flow cytometry, and the percentages of apoptotic cells are shown (C). Apoptosis-related protein levels were analyzed by western blot, with densitometric quantification relative to β-actin shown (D). (E and F) c-Myc-knockdown A549 cells were reconstituted with Flag-tagged wild-type c-Myc and then mock-treated or infected with NDV at an MOI of 1 for 24 h. Apoptosis was analyzed by flow cytometry, and the percentages of apoptotic cells are shown (E). Apoptosis-related protein levels were analyzed by western blot, with densitometric quantification relative to β-actin shown (F). (G and H) c-Myc-knockdown A549 cells were reconstituted with Flag-tagged c-Myc K142R and then mock-treated or infected with NDV at an MOI of 1 for 24 h. Apoptosis was analyzed by flow cytometry, and the percentages of apoptotic cells are shown (G). Apoptosis-related protein levels were analyzed by western blot, with densitometric quantification relative to β-actin shown (H). Data are presented as means from three independent experiments. *P < 0.05, ***P < 0.001; ns, not significant.

## DISCUSSION

As a transcription factor, c-Myc plays a critical role in regulating cell proliferation and differentiation. However, aberrant c-Myc expression is frequently observed in tumors, where it functions as a well-established oncogenic driver (17, 18). Due to the lack of a well-defined binding pocket, c-Myc has long been considered difficult to target directly with small molecules, and clinically effective c-Myc-targeted therapies remain limited (20, 32). Recent studies have explored several indirect strategies to suppress c-Myc activity. For example, Tasquinimod-mediated inhibition of the HDAC4–c-Myc axis has been proposed as a potential therapeutic strategy for multiple myeloma (33). In addition, cell-permeable small-molecule MYC inhibitors have been developed to disrupt MYC/MAX dimerization, enhance MYC phosphorylation at threonine 58, and promote proteasome-mediated MYC degradation, thereby suppressing tumor growth and improving antitumor immune responses in preclinical models (34). Another small-molecule inhibitor, 5D4, suppresses MYC activity by disrupting the TopBP1–MIZ1 interaction and restoring MIZ1 binding to target gene promoters (35). Despite these advances, the therapeutic efficacy of c-Myc-targeted strategies remains to be fully established. Here, we demonstrate that oncolytic NDV promotes SUMO-1-dependent nuclear translocation of PRKN, enabling PRKN to interact with c-Myc and mediate its ubiquitination-dependent degradation at Lys142. This process activates apoptotic signaling and suppresses tumor growth, revealing a distinct mechanism by which NDV targets the difficult-to-drug oncogenic driver c-Myc (Fig. 8).

**Figure 8.**
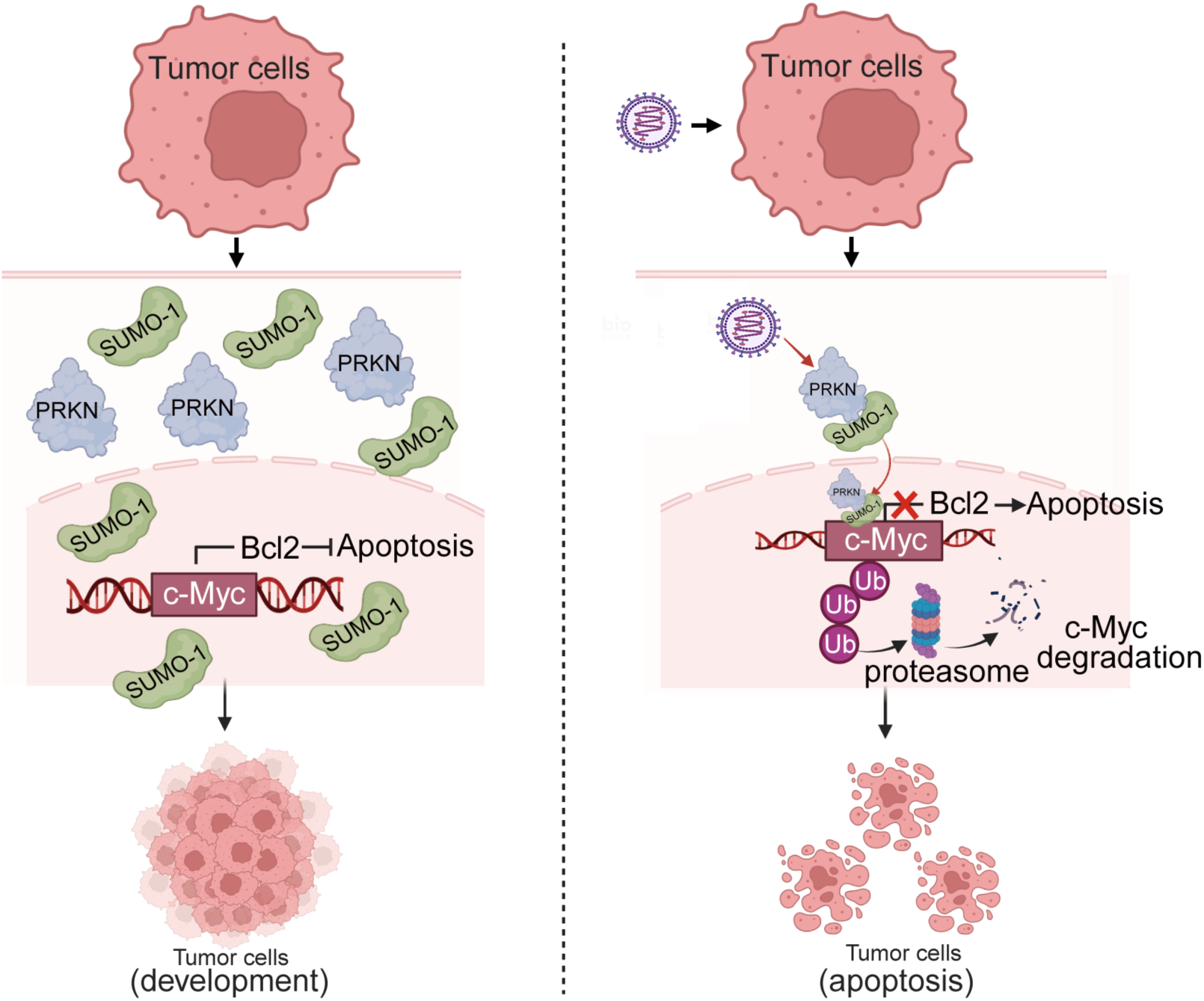
Proposed model of NDV-induced c-Myc degradation and tumor cell apoptosis. Schematic model showing that NDV infection promotes SUMO-1-dependent nuclear translocation of PRKN. Nuclear PRKN mediates ubiquitination of c-Myc at Lys142, leading to proteasomal degradation of c-Myc. Loss of c-Myc decreases Bcl-2 expression and activates the intrinsic apoptotic pathway, thereby promoting tumor cell apoptosis.

Notably, NDV-induced c-Myc degradation was observed across multiple tumor cell types and was induced by NDV strains of different pathotypes, including velogenic, mesogenic and lentogenic strains. This is consistent with previous studies showing that NDV displays broad oncolytic activity in diverse tumor models, largely due to defects in antiviral defense pathways, particularly impaired type I IFN signaling, in cancer cells (36–39) Although NDV strains differ in virulence, replication capacity and fusogenic activity, both lytic and less lytic strains have been reported to activate tumor cell death and antitumor immune responses (40). Therefore, the ability of different NDV pathotypes to reduce c-Myc levels suggests that c-Myc destabilization is not restricted to a specific viral strain, but may represent a relatively conserved host response to productive NDV infection. In contrast, UV-inactivated NDV failed to induce c-Myc degradation, indicating that active viral infection, viral replication or de novo viral gene expression is required for this process. This observation is in line with previous reports showing that NDV-induced apoptosis depends on viral entry, replication and de novo protein synthesis (40, 41). The extent of NDV-induced apoptosis differed among tumor cell lines, which may reflect tumor-intrinsic differences in antiviral signaling competence, viral receptor availability, apoptotic threshold, and expression of anti-apoptotic factors such as Bcl-2 family proteins or inhibitor of apoptosis proteins (36, 42). In the context of our findings, basal c-Myc abundance, PRKN expression and SUMO-1-dependent PRKN nuclear translocation may further influence cellular sensitivity to NDV. Thus, c-Myc expression and the activity of the SUMO-1-PRKN axis, together with established determinants such as IFN pathway status, may serve as potential biomarkers for predicting tumor responsiveness to NDV-based therapy.

NDV-mediated oncolysis is not restricted to apoptosis but involves multiple forms of regulated cell death. Apoptosis is a well-established mechanism of NDV-induced tumor cell killing, involving mitochondrial dysfunction, Bax/Bcl-2 imbalance, caspase activation, and death receptor-related signaling (36, 43, 44). NDV can also activate ER stress-, UPR- and MAPK-related pathways that contribute to apoptotic cell death (45, 46). Beyond apoptosis, NDV has been reported to induce necroptosis, immunogenic cell death, autophagy-dependent immunogenic cell death, ferroptosis, parthanatos and anoikis in different tumor contexts (4, 5, 8, 47–51). These findings indicate that NDV activates a complex network of tumor cell death programs. Our study adds c-Myc ubiquitination and degradation as a new mechanism underlying NDV-mediated apoptosis. Given the broad role of c-Myc in regulating mitochondrial metabolism, redox homeostasis, amino acid metabolism, iron metabolism and survival signaling, future studies should determine whether the NDV-SUMO-1-PRKN-c-Myc axis also contributes to other regulated cell death pathways, such as ferroptosis, parthanatos, autophagy-associated cell death and anoikis.

Several E3 ubiquitin ligases have been reported to regulate c-Myc stability. For example, FBXL14 promotes c-Myc degradation, induces glioma stem cell differentiation, and suppresses tumor growth (52). FBW7, another well-characterized E3 ubiquitin ligase, reduces c-Myc protein stability by promoting its phosphorylation-dependent ubiquitination and subsequent proteasomal degradation (53). In contrast to these previously described mechanisms, our study identifies Lys142 as a critical residue involved in NDV-induced ubiquitination-dependent degradation of c-Myc. This suggests that NDV activates a distinct c-Myc destabilization pathway that differs from the classical phosphorylation-dependent Thr58–FBW7 axis. PRKN is predominantly localized in the cytoplasm and plays important roles in cellular homeostasis, mitophagy, and neuroprotection (54). However, our results showed that PRKN translocates into the nucleus following NDV infection, where it mediates c-Myc ubiquitination and degradation. This finding expands the functional scope of PRKN beyond its canonical role in mitochondrial quality control and suggests that PRKN may acquire noncanonical nuclear functions during viral infection.

An important question raised by our findings is how PRKN nuclear translocation is regulated during NDV infection. Previous studies have shown that PRKN selectively associates with SUMO-1 both in vivo and in vitro, and that SUMO-1 binding promotes PRKN nuclear transport and enhances its autoubiquitination activity (31). SUMOylation and SUMO-related regulation play pleiotropic roles during viral infection by modulating protein–protein interactions and directing the subcellular or subnuclear localization of target proteins (55). onsistently, our study found that SUMO-1 knockdown inhibited NDV-induced nuclear translocation of PRKN, thereby blocking PRKN-mediated c-Myc ubiquitination and degradation and attenuating NDV-induced tumor cell apoptosis. These findings suggest that SUMO-1 acts as a key regulator of PRKN subcellular trafficking during NDV infection. However, whether NDV directly promotes SUMOylation of PRKN, and which viral component or infection-associated signal initiates SUMO-1-dependent PRKN nuclear translocation, remain to be further investigated.

In conclusion, our study uncovers a novel mechanism by which NDV exerts its oncolytic activity. We demonstrate that NDV activates a SUMO-1-PRKN-c-Myc regulatory axis, in which SUMO-1 facilitates the nuclear translocation of PRKN and allows PRKN to ubiquitinate and degrade c-Myc at Lys142. By destabilizing c-Myc, NDV promotes tumor cell apoptosis and suppresses tumor growth. These findings not only expand the mechanistic understanding of NDV-mediated antitumor activity but also highlight c-Myc destabilization as a potential therapeutic strategy for cancer treatment.

## MATERIALS AND METHODS

### Antibodies and Reagents

The following primary antibodies were used for western blotting: anti-β-actin (Proteintech, Cat# 66009-1-Ig), anti-NDV NP antibody generated in our laboratory, anti-PARP (Cell Signaling Technology, CST, Cat# 9542S), anti-cleaved caspase-3 (CST, Cat# 9661S), anti-caspase-3 (CST, Cat# 9662S), anti-caspase-8 (CST, Cat# 4790S), anti-cleaved caspase-8 (CST, Cat# 9496S), anti-HA (CST, Cat# 3724S), anti-β-tubulin (CST, Cat# 5668S), anti-c-Myc (Proteintech, Cat# 67447-1-Ig), anti-SOX2 (Proteintech, Cat# 11064-1-AP), anti-KLF4 (Proteintech, Cat# 11880-1-AP), anti-OCT4 (Proteintech, Cat# 11263-1-AP), anti-Bcl-2 (Proteintech, Cat# 12789-1-AP), anti-caspase-9 (CST, Cat# 9502S), anti-LC3 (Proteintech, Cat# 14600-1-AP), anti-Flag (Sigma-Aldrich, Cat# F1804), anti-PRKN (Proteintech, Cat# 14060-1-AP), anti-Lamin B1 (Proteintech, Cat# 12987-1-AP), and anti-SUMO-1 (Proteintech, Cat# 10329-1-AP). Alexa Fluor 488-conjugated secondary antibody (Thermo Fisher Scientific, Cat# A-11008) and Alexa Fluor 594-conjugated secondary antibody (Thermo Fisher Scientific, Cat# A-11037) were used for immunofluorescence staining.

The Annexin V-FITC Apoptosis Detection Kit (Cat# C1062L) and DAPI (Cat# C1002) were purchased from Beyotime. MG132 (Cat# S2619) was purchased from Selleck Chemicals. Chloroquine (CQ, Cat# C6628) was purchased from Sigma-Aldrich. Pierce Protein A/G Magnetic Beads (Cat# 88847) were purchased from Thermo Fisher Scientific.

### Plasmids

Flag-tagged c-Myc (Flag-c-Myc) and HA-tagged c-Myc (HA-c-Myc) were generated by inserting the ORF of human c-Myc (NM_002467.6) into the p3XFLAG-CMV-14 vector (Sigma-Aldrich) and pCMV-HA vector (Clontech), respectively. Flag-tagged c-Myc point mutants, including K67R, K68R, K67/68R, K142R, K159R, K164R, K173R, K222R, K285R, K291R were generated using the Mut Express II Fast Mutagenesis Kit V2 (Vazyme, China) according to the manufacturer’s instructions. Flag-tagged PRKN (Flag-PRKN) was generated by inserting the ORF of human PRKN (NM_004562.3) into the p3XFLAG-CMV-14 vector. pRK5-HA-ubiquitin-WT was obtained from Addgene (Cat#17608).

### Cell culture, siRNA transfection, and viruses

A549, HeLa, MCF-7, PANC-1, and HT-29 cells were purchased from the Cell Bank of the Shanghai Institute of Biochemistry and Cell Biology, Chinese Academy of Sciences. A549 cells were cultured in RPMI-1640 medium, whereas HeLa, MCF-7, PANC-1, and HT-29 cells were cultured in Dulbecco’s modified Eagle’s medium (DMEM). All media were supplemented with 10% fetal bovine serum (FBS; ExCell Bio, Cat# FSP500), and cells were maintained at 37°C in a humidified incubator with 5% CO₂. Antibiotics were not added to the culture medium during the experiments.

Gene silencing was performed by transfecting cells with siRNAs using Lip ofectamine 2000 in Opti-MEM reduced-serum medium according to the manufacturer’s instructions. The siRNAs were used at 100 pmol per well unless other wise indicated. The following siRNA sequences were used: si-c-Myc #1, 5′-GA GGAUAUCUGGAAGAAAUTT-3′ (42); siTRPC4AP, 5′-GUGGAAUUUGUGGA ACAUCTT-3′; siSTUB1, 5′-GGCAAUCGUCUGUUCGUGGGCCGAA-3′; siFBX W7, 5′-CACAAAGCUGGUGUGUGCATT-3′; siRNF187, 5′-CACUGAGCGGUU CAGGUCATT-3′; siPRKN, 5′-CCUUCUGCCGGGAAUGUAAAGAAGCGUAC-3′; siFBXW11, 5′-GGUUGUUAGUGGAUCAUCATT-3′; siSKP2, 5′-CCUAUCGAA CUCAGUUAUATT-3′ (23); and siSUMO-1, 5′-GGACAGGATAGCAGTGAGA-3′ (43). All siRNAs were synthesized by GenePharma (Shanghai, China).

The NDV strains Herts/33 and La Sota were obtained from the China Institute of Veterinary Drug Control (Beijing, China), and the Mukteswar strain wa s obtained from Yangzhou University (Yangzhou, China). Unless otherwise state d, Herts/33 was used as the default NDV strain in this study. UV-inactivated NDV was prepared by exposing the Herts/33 virus stock to ultraviolet irradiation for 3 h. For infection, cells were incubated with NDV at a multiplicity of infection (MOI) of 1, unless otherwise indicated, for 1 h at 37°C. The inoculum was then removed and replaced with antibiotic-free maintenance medium con taining 1% FBS. Where indicated, inhibitors were added to the culture medium and cells were harvested at the specified time points. Extracellular NDV titers were determined in DF-1 cells as the 50% tissue culture infectious dose (TCI D_50_) using the Reed-Muench method.

### Western blot analysis

Cells cultured in 6-well plates were harvested, washed with PBS, and lysed directly in 2× loading buffer containing 20 mM Tris-HCl, 2% SDS, 100 mM dithiothreitol, 20% glycerol, and 0.016% bromophenol blue. Samples were boiled for 15 min and stored at −20°C until use. Proteins were separated by 10% SDS-PAGE and transferred onto nitrocellulose membranes (Whatman, Cat# NC-a101b105). After blocking with 5% skimmed milk for 1 h at room temperature, the membranes were incubated with primary antibodies overnight at 4°C. After washing, the membranes were incubated with HRP-conjugated secondary antibodies for 2 h at room temperature. Protein bands were detected using a basic ECL chemiluminescence substrate (Share-bio, Shanghai, China) and imaged with a chemiluminescence detection system (Tanon, Shanghai, China).

### Co-immunoprecipitation

HEK-293T cells were lysed on ice for 30 min in RIPA lysis buffer supplemented with protease inhibitor cocktail. Cell lysates were incubated with anti-Flag magnetic beads (Bimake, Houston, TX, USA) or anti-HA magnetic beads (Share-Bio, Shanghai, China) overnight at 4°C. The beads were washed three times with wash buffer containing 150 mM NaCl, 50 mM Tris-HCl, pH 8.0, 5 mM EDTA, and 0.5% NP-40. Bound proteins were eluted by boiling in SDS loading buffer containing 10 mM Tris-HCl, pH 7.8, 3% SDS, 5% glycerol, and 0.02% bromophenol blue for 10 min, and then analyzed by western blot.

### Apoptosis assay

Apoptosis was assessed using an Annexin V-FITC Apoptosis Detection Kit according to the manufacturer’s instructions. A549, HeLa, MCF-7, PANC-1, and HT-29 cells were seeded in 6-well plates at 1 × 10⁶ cells per well and infected with NDV as indicated. Cells were harvested, stained with Annexin V-FITC and propidium iodide (PI), and analyzed by flow cytometry.

### Animal studies

Six-week-old BALB/c nude mice, with an average body weight of 18 g, were purchased from Shanghai Jiesijie Laboratory Animal Co., Ltd. Mice were housed under standard conditions with aspen chip bedding, in accordance with institutional guidelines. For the subcutaneous xenograft model, A549 cells, 1 × 10^7^ cells per mouse, were suspended in cold DMEM and subcutaneously injected into the rear flank of each mouse. Tumor size was measured using digital calipers, and tumor volume was calculated using the following formula: tumor volume, mm³ = 0.52 × length × width^2^. Seven days after tumor cell implantation, mice were randomly divided into the indicated groups. To evaluate the antitumor effect of NDV, mice were intravenously injected with NDV, 10^8.5^ EID_50_/100 μL, or saline as a negative control twice per week for approximately 24 days. Tumor size and body weight were recorded twice weekly. All animal experiments were conducted according to protocols approved by the

Animal Care and Use Committee of the Chinese Academy of Agricultural Sciences (CAAS; ethical approval number SV-20231027-G01).

### Immunofluorescence staining

Cells grown on glass coverslips were washed once with PBS and fixed with 4% paraformaldehyde for 30 min at room temperature. Cells were then permeabilized with 0.5% Triton X-100 for 10 min and blocked with 5% bovine serum albumin (BSA) for 2 h at room temperature. Samples were incubated with the indicated primary antibodies overnight at 4°C, washed three times with PBST, and then incubated with the appropriate secondary antibodies for 2 h at room temperature. After nuclear staining with DAPI, images were acquired using an LSM880 confocal microscope (Carl Zeiss, Jena, Germany).

### Statistical analysis

Data are presented as the mean ± SD. Fluorescence intensities and western blot band intensities were quantified using ImageJ software. Statistical analyses were performed using GraphPad Prism. Comparisons between two groups were performed using an unpaired Student’s t-test, whereas comparisons among multiple groups were analyzed by one-way ANOVA. A value of P < 0.05 was considered statistically significant.

## DATA AVAILABILITY

The data generated in this study are included in this article. Additional data are available from the corresponding author upon reasonable request.

## AUTHORS’ CONTRIBUTIONS

Changrun Zhao: Validation, Visualization, Methodology, Writing – original draft. Ning Tang: Visualization, Methodology. Liya Shen: Visualization, Methodology. Jing Wang: Visualization. Sonshu Meng: Validation. Jianghong Yu: Validation. Yiqian Zhu: Validation. Zunping Ke: Validation. Yang Qu: Validation. Lei Tan: Validation. Cuiping Song: Validation. Xusheng Qiu: Validation. Ying Liao: Validation. Yingjie Sun: Supervision, Funding acquisition. Chan Ding: Supervision, Funding acquisition, Investigation.

## ACKNOWLEDGMENTS

This study was supported by the National Natural Science Foundation of China (32530102), the International Cooperation Project of National Natural Science Foundation of China (32220103012), National Natural Science Foundation of China (32503020) and the Innovation Program of the Chinese Academy of Agricultural Sciences (CAAS-CSLPDCP-202402).

